# Neuromuscular fatigue and recovery after strenuous exercise depends on skeletal muscle size and stem cell characteristics

**DOI:** 10.1101/740266

**Authors:** P. Baumert, S. Temple, J.M. Stanley, M. Cocks, J.A. Strauss, S.O. Shepherd, B. Drust, M.J. Lake, C.E. Stewart, R.M. Erskine

## Abstract

Hamstring muscle injury is highly prevalent in sports involving repeated maximal sprinting. Although neuromuscular fatigue is thought to be a risk factor, the mechanisms underlying the fatigue response to repeated maximal sprints are unclear. Here, we show that repeated maximal sprints induce neuromuscular fatigue accompanied with a prolonged strength loss in hamstring muscles. The immediate hamstring strength loss was linked to both central and peripheral fatigue, while prolonged strength loss was associated with indicators of muscle damage. The kinematic changes immediately after sprinting likely protected fatigued hamstrings from excess elongation stress, while larger hamstring muscle physiological cross-sectional area and lower *myoblast:fibroblast* ratio appeared to protect against fatigue/damage and improve muscle recovery within the first 48 h after sprinting. We have therefore identified novel mechanisms that likely regulate the fatigue/damage response and initial recovery following repeated maximal sprinting in humans.

## INTRODUCTION

Hamstring strain is the most frequently occurring injury in sport ^1^, particularly in those sports that involve high-speed running ^2^. Although the aetiology is unclear, numerous risk factors have been proposed, such as short fascicle length, poor flexibility, poor hamstring strength, and inadequate warm-up ^3^. Further, it is unknown whether hamstring strain is the result of a single event that exceeds the physiological range of hamstring muscle extensibility and contractility, or as a result of an accumulation of eccentric contractions during repeated maximal sprints, causing neuromuscular fatigue ^3^. Neuromuscular fatigue is responsible for acute, as well as prolonged, impairment of muscle function, classified as central fatigue (i.e. originating in the central nervous system), or peripheral fatigue (i.e. distal to the neuromuscular junction) ^4^. Although it was recently reported that both central and peripheral fatigue contribute to impaired hamstring muscle function immediately after repeated maximal sprint-related interventions ^5,6^, the contribution of neuromuscular fatigue to hamstring muscle impairment and recovery, following repeated maximal sprints over time, is insufficiently studied ^7^. An understanding of hamstring neuromuscular fatigue following repeated maximal sprints may be crucial for understanding hamstring strain aetiology.

Peripheral fatigue may be caused by ultrastructural muscle damage, which is indicated by Z-line disturbance ^8^ as well as disruption of the extracellular matrix ^9^. The extracellular matrix provides structural scaffolding for muscle remodelling and plays an integral role in force transmission ^10^. This is referred to as exercise-induced muscle damage and it is exhibited by prolonged strength loss and delayed-onset muscle soreness, as well as the release of muscle-specific proteins [e.g. creatine kinase (CK)] into the circulation over the following days ^11^. After substantial muscle damage, myogenic satellite cells (skeletal muscle stem cells), play a key role in skeletal muscle regeneration and remodelling ^12^. Activated satellite cells (myoblasts) proliferate and migrate from their niche along the basal lamina to the injury site before terminally differentiating and fusing with damaged myofibrils to repair injury. There is increasing evidence that fibroblasts, the main cell type of muscle connective tissue, also play a critical role in supporting muscle regeneration ^13,14^. Following damage, infiltrating inflammatory cells activate muscle fibroblasts, which proliferate and migrate to the area of the myotrauma and produce extracellular matrix components in an orchestrated and regulated fashion to support healthy muscle remodelling ^14,15^. However, little is known about the role of fibroblasts during the initial response and recovery following *physiological* exercise-induced muscle damage, e.g. following repeated sprinting.

Acute damage to the muscle-tendon complex may facilitate hamstring strain, which is thought to occur in the late swing phase of sprinting, when the hamstring muscles contract eccentrically, i.e. trying to shorten while being lengthened in an attempt to decelerate the shaft before initial foot-ground contact ^16^. Therefore, a short biceps femoris long head (BF_LH_) fascicle length has been suggested to increase hamstring strain risk ^17^, as the BF_LH_ is thought to be relatively more eccentrically stretched during the late swing phase of sprinting compared to the other hamstring muscles ^16^. However, no study has investigated the relationship between BF_LH_ architecture (including muscle fascicle length and cross-sectional area), and the prolonged hamstring muscle response to exercise-induced neuromuscular fatigue. Finally, lower limb neuromuscular fatigue might cause a number of biomechanical alterations in running kinematics ^18^. However, it is not known whether repeated maximal sprints influence kinematic patterns, and whether this can lead to prolonged changes in lower-limb kinematics, which may play a role in the development of muscle strain following insufficient recovery ^3^.

Here we demonstrated that both central and peripheral fatigue, assessed via surface electromyography (sEMG) and electrical stimulation, contributed to the immediate loss of muscle function in both the quadriceps and hamstring muscle groups, but that peripheral factors mainly contributed to the sustained loss of hamstring muscle function. Moreover, we established that a lower myoblast:fibroblast ratio in isolated primary human muscle stem cells correlated with improved recovery from both repeated maximal sprints and an *in vitro* artificial wounding assay within the first 48 h. We also report that BF_LH_ architecture (i.e. physiological cross-sectional area, PCSA) was associated with hamstring fatigue, and that neuromuscular fatigue led to reduced hip flexion and knee extension during the late swing phase of steady-state running. Thus, with this interdisciplinary study, we identify novel cellular and neuromuscular mechanisms underpinning central and peripheral fatigue following repeated sprinting, which ultimately led to kinematic changes during the running stride phase associated with hamstring strain injury.

## RESULTS

### Effect of the Repeated Maximal Sprint Intervention on Neuromuscular Fatigue

To examine the effect of repeated maximal sprints on neuromuscular fatigue, we measured different fatigue parameters before (PRE), immediately after (POST), and 48 h after (POST48) the repeated maximal sprint intervention. The average 30 m sprinting speed was 6.48 ± 0.33 m s^−1^. There was a main effect of time for heart rate, 30 m sprinting time, rating of perceived exertion and lactate concentration, with all parameters increasing from PRE to POST (all P<0.001). Blood lactate concentration increased from PRE (1.63 ± 0.45 mmol/L) to POST (9.82 ± 3.62 mmol/L; P<0.001). The sprinting performance (measured via the performance decrement score ^19^) decreased by 3.98 ± 2.99% during the run, and rating of perceived exertion increased by 96.5 ± 35.2% from PRE-to-POST, indicating fatigue had occurred.

We then performed *in vivo* functional analysis to assess if repeated maximal sprints resulted in an increase in central and/or peripheral fatigue. We, therefore, measured BF_LH_ muscle activation via normalised surface electromyography (sEMG) during hamstring maximum voluntary contraction (MVC). We observed a change in sEMG (FF_2,24_=4.35, P=0.022), with post-hoc pairwise comparisons revealing a decrease from PRE-to-POST (−24.3%; P=0.019). However, this change was no longer evident at POST48 (P=0.157, Table 1), suggesting that central fatigue occurred immediately after repeated maximal sprints. No other changes in muscle (co)activation were observed at any time point (P>0.05).

**Table 1.**
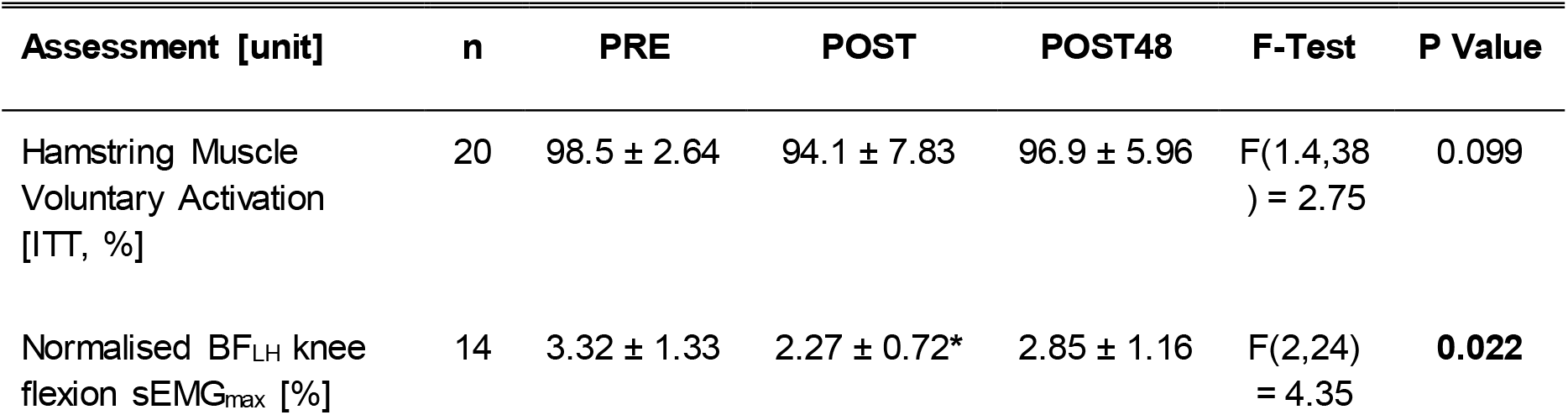
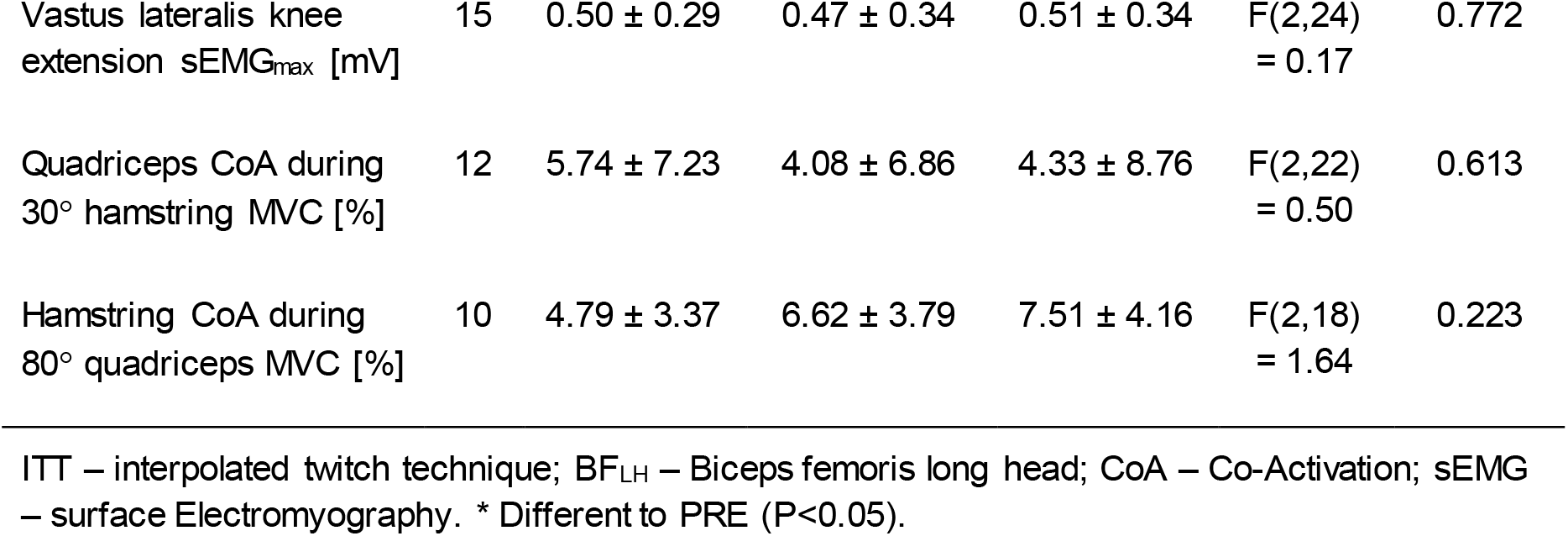
Effect of the repeated maximal sprint intervention on measures of muscle activation. Data are presented as mean ± SD. One-way ANOVA, F- and P-values are reported.

We also assessed the torque-frequency relationship *in vivo* via electrical stimulation to indicate peripheral (muscle) fatigue. There was an interaction between time x stimulation frequency (n=19; F_4.9,216_=6.62, P<0.001; Figure 1A). Post-hoc paired t-tests revealed differences PRE-to-POST for 10-50 Hz (P<0.05), but lower frequencies between 10-20 Hz reverted to baseline values POST48 (P>0.05), while the frequencies of 30 and 50 Hz were still decreased POST48 compared to their baseline values (P<0.05), providing evidence that peripheral fatigue occurred immediately after the repeated sprints and remained for 48 h

**Figure 1.**
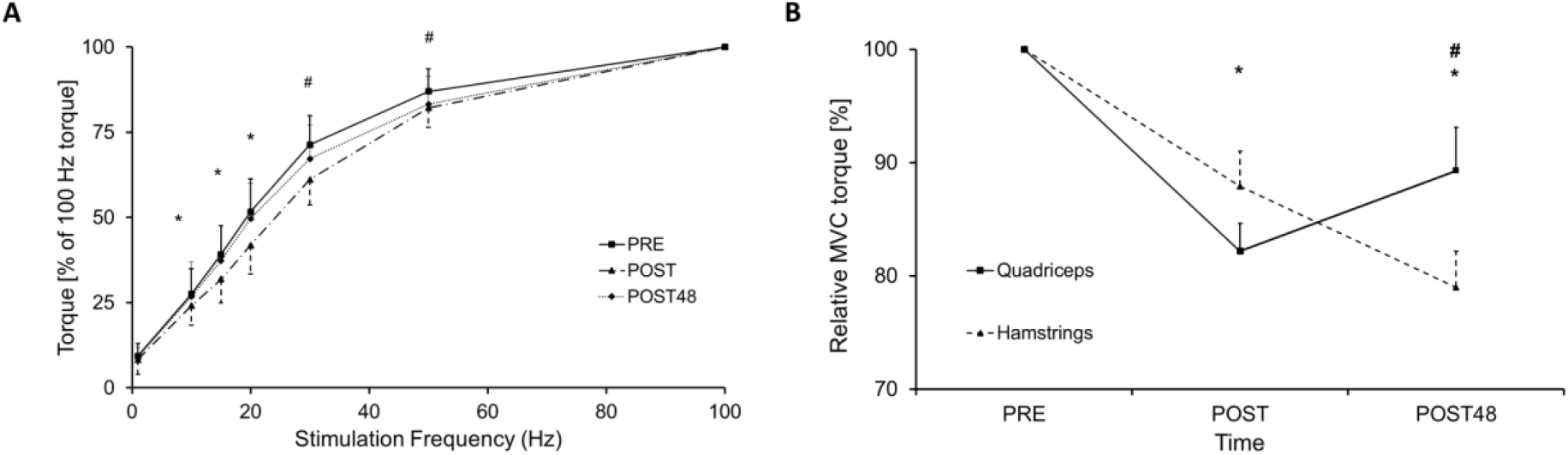
**(A)** Torque-frequency relationship, all frequencies were normalised to 100 Hz. * significant differences between before (PRE) and immediadtely after (POST) the repeated maximal sprint intervention, P<0.05; # significant differences between PRE and POST, and between PRE and POST48, P<0.05. Data are presented as mean ± SEM. **(B)** Comparison of relative maximal voluntary contraction (MVC) loss between hamstring and quadriceps muscle group before (PRE), immediately after (POST) and 48h after (POST48) the repeated maximal sprint intervention. * signifcant differences compared to PRE, P< 0.001; # significant differences between quadriceps and hamstring MVC, P< 0.05. Data are expressed as mean ± SEM.

### Effect of the Repeated Maximal Sprint Intervention on MVC Strength, Muscle Soreness and Serum Markers of Exercise-Induced Muscle Damage

To investigate the effect of repeated maximal sprints on biomarkers of exercise-induced muscle damage, we assessed hamstring (knee flexion) and quadriceps (knee extension) MVC, muscle soreness, serum creatine kinase (CK) activity and interleukin-6 (IL-6) concentrations PRE, POST and POST48. Isometric hamstring and quadriceps MVC, muscle soreness (all P<0.001) and serum CK activity (F_1.3,34_=5.98, P=0.017), as well as IL-6 concentration (F_1.3,34_=5.96, P=0.018), showed a main effect of time, which are indicators of muscle damage) that was similar to other studies ^7,20,21^. Post-hoc pairwise comparisons revealed that, compared to PRE, both serum CK activity (+93.0%) and serum IL-6 concentration (+307%) were elevated at POST (both P=0.027), and CK activity further increased at POST48 (+256%; P=0.012), while serum IL-6 concentration reverted to baseline values (P>0.05).

Further, there was an interaction between time and muscle groups concerning relative MVC torque loss (percentage change from PRE MVC) (F_1.4,38_=7.92, P=0.004). Relative MVC decreased similarly in both quadriceps and hamstring muscle groups PRE-to-POST (Figure 1B). However, at POST48, hamstring MVC continued to decrease from POST (−9.26%; P=0.010), while quadriceps MVC began to return to PRE values (+8.47%; P=0.016) and was higher than hamstring MVC at POST48 (P=0.038).

### Effect of the Repeated Maximal Sprint Intervention on Lower-Limb Kinematics

To assess the consequential effect of neuromuscular fatigue on lower-limb kinematics, we captured treadmill running (4.17 m s^−1^) kinematics with an eight-camera motion capture system over time. Three-dimensional motion analysis demonstrated, that despite significance not being achieved, there was a tendency towards a longer running cycle time POST (+1.01%) and POST48 (+1.86%), compared to PRE (P=0.080; Table 3). Further, treadmill running demonstrated decreased peak knee extension (P=0.047) during the late swing phase at POST (−10.9%) compared to PRE, but this reverted to baseline POST48. The percentage change in peak knee extension correlated with the percentage change in relative hamstring MVC torque both measured POST-to-POST48 (R^2^=0.26, F_1,2_=5.673, P=0.031).

**Table 2.**
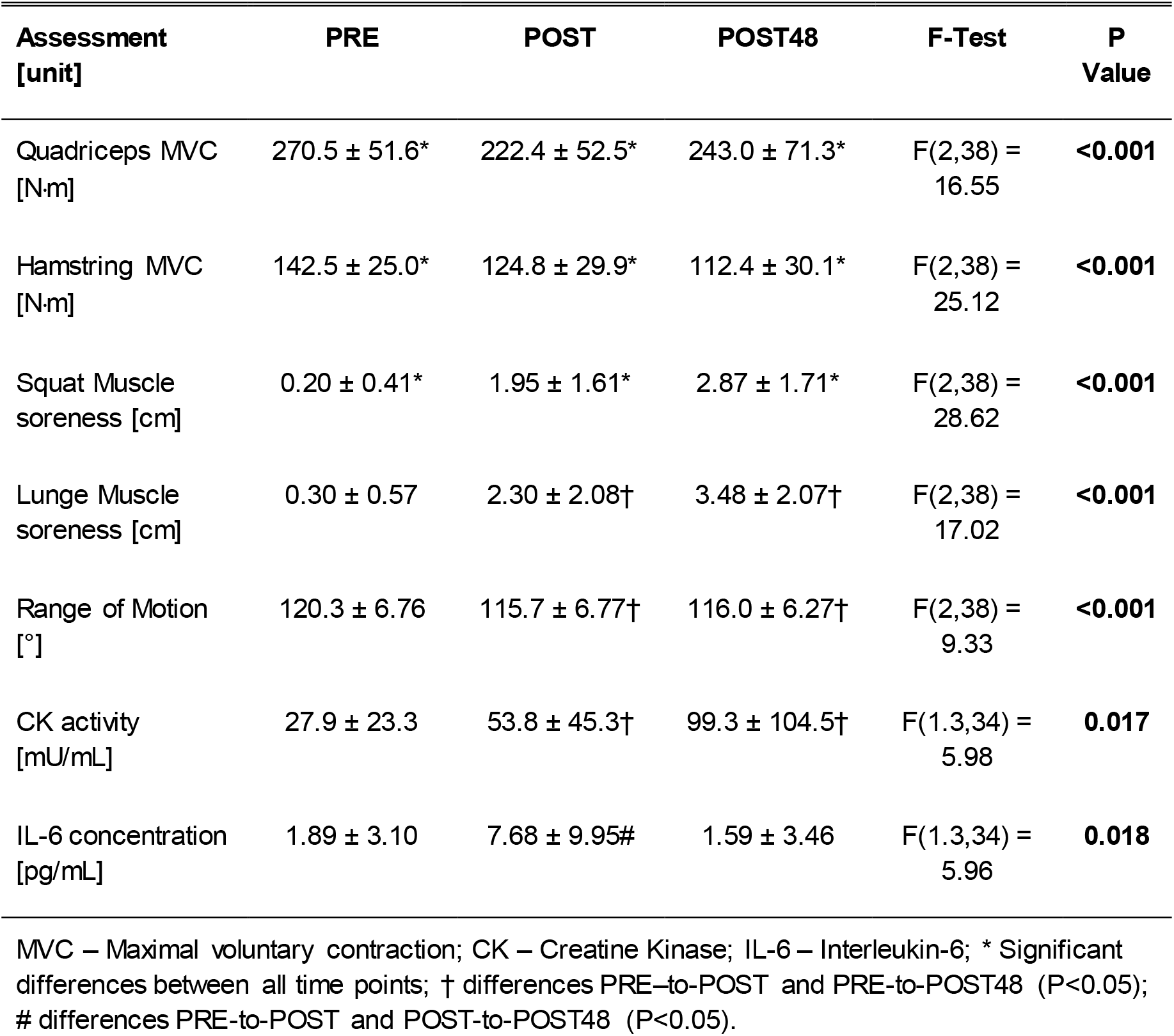
Effect of the Repeated Maximal Sprint Intervention on Muscle Damage-Biomarkers. Values are mean ± SD. One-way ANOVA, F- and P-values are reported.

**Table 3.** Effect of the Repeated Maximal Sprint Intervention on Kinematics of Treadmill Running at 4.17 m s−1 Values are mean ± SD. One-way ANOVA, F- and P-values are reported.

| Kinematics [unit] | PRE | POST | POST48 | F-Test | P Value |
| --- | --- | --- | --- | --- | --- |
| Peak knee flexion (swing phase) [°] | -103 ± 13.9 | -110 ± 12.4 | -108 ± 12.6 | F(2,20) = 2.84 | 0.082 |
| Peak knee extension (swing phase) [°] | -3.66 ± 5.32 | -7.08 ± 5.07* | -4.29 ± 7.06 | F(2,20) = 3.57 | **0.047** |
| Contact hip flexion (toe strike) [°] | 25.2 ± 5.42 | 29.3 ± 7.62 | 17.6 ± 15.6 | F(1.2,20) = 4.05 | 0.062 |
| Contact knee flexion (toe strike) [°] | -15.0 ± 6.25 | -17.6 ± 6.79 | -13.3 ± 9.64 | F(2,22) = 2.79 | 0.083 |
| Duration Running cycle [s] | 0.67 ± 0.03 | 0.68 ± 0.03 | 0.69 ± 0.02 | F(2,20) = 2.88 | 0.080 |
| Stance phase duration [s] | 0.18 ± 0.02 | 0.19 ± 0.02 | 0.19 ± 0.03 | F(2,20) = 1.01 | 0.356 |
| Swing Phase [s] | 0.50 ± 0.04 | 0.50 ± 0.05 | 0.50 ± 0.03 | F(1.2,20) = 0.03 | 0.899 |

Knee fully extended=0°; negative value indicates a flexed knee; \* different to PRE (P<0.05).

### Architecture of the Biceps Femoris Long Head Muscle

To assess whether architectural parameters of the BF_LH_ muscle (Figure 2) were associated with markers of peripheral fatigue, we performed ultrasound measurements of the BF_LH_ muscle (Table supplement 4). Muscle fascicle length and pennation angle of the BF_LH_, which have previously been linked to hamstring muscle strain risk ^22^, did not correlate with any outcome variable of neuromuscular fatigue. However, BF_LH_ PCSA (mean ±SD: 23.4 ± 4.62 cm^2^) correlated inversely with relative hamstring MVC loss PRE-to-POST (R^2^=0.42, F_1,17_=12.37, P=0.003, Figure 2C).

**Figure 2.**
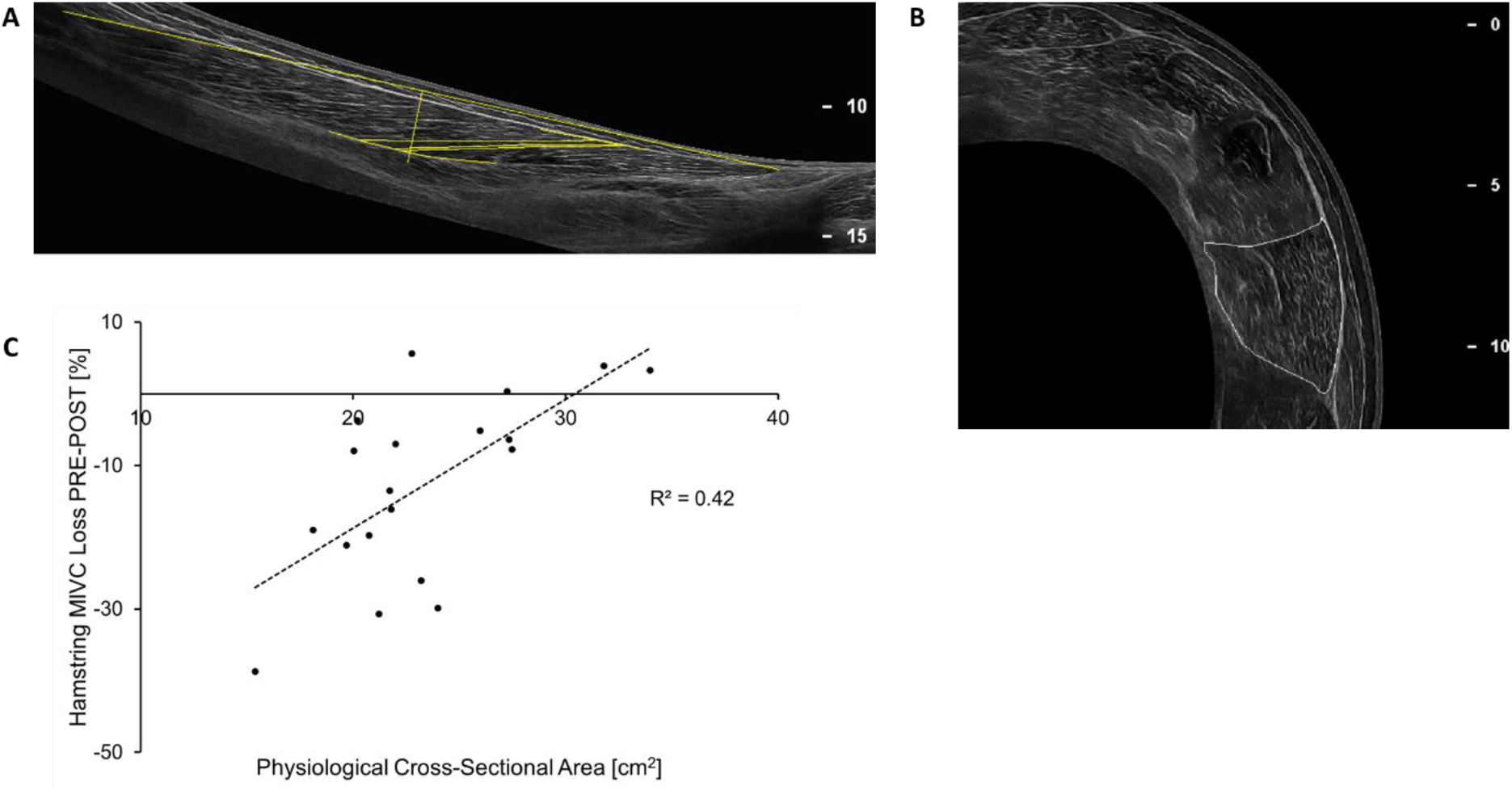
**(A)** Longitudinal image of biceps femoris long head, assessment of the biceps femoris long head is highlighted (total muscle length and fascicle length together with pennation angle at 50% of total muscle length). **(B)** Cross-sectional image at 60% muscle length (=100% proximal myotendinous junction), biceps femoris long head is highlighted. **(C)** Correlation between biceps femoris long head muscle physiological cross-sectional area and % hamstring maximum isometric voluntary contraction (MIVC) decrease from before (PRE) to immediately after (POST) the repeated maximal sprint intervention (P=0.003).

### Artificial Wound Healing Assay to Investigate Repair and Regeneration Regarding Myoblast:Fibroblast Ratio

Preliminary data from our laboratory demonstrated that skeletal muscle stem cell composition (i.e. myoblast:fibroblast ratio), derived from two volunteers with ratios representing the extreme conditions of the myoblast:fibroblast percentage, played a role in the success of artificial wound healing *in vitro* ^23^. Human primary skeletal muscle stem cells with a high myoblast:fibroblast ratio resulted in reduced cell migration into an artificial wound (assessed by cell number within the wound) when compared with stem cells with a low myoblast:fibroblast ratio. No significant differences were observed in the relative proportion of migrating cells (i.e. between myoblasts and fibroblasts) over 48 hours. Further, recent investigations have revealed that the skeletal muscle stem cell ratio does not change during *in vitro* cell culturing ^24,25^. We, therefore, further assessed the effect of the myoblast:fibroblast ratio on skeletal muscle recovery following *in vitro* artificial wounding to extend the preliminary *in vitro* results from our laboratory. To be able to demonstrate a coefficient of determination of ≥ 0.50 between myoblast:fibroblast ratio and our dependent variables, a priori power calculations for a Pearson's r correlation was run using G*Power (version 3.1.9.1), and it revealed that 11 persons provided alpha = 5%, and power = 80%. Thus, we used primary human skeletal muscle stem cells derived from 12 participants, six who participated in both the repeated maximal sprint intervention and also volunteered to provide a muscle biopsy at least three weeks before the repeated maximal sprint intervention, and another six (two male and four females), who did not participate in the repeated maximal sprint intervention (to increase the power of the *in vitro* study). The muscle stem cells were isolated, cultured and then characterized by immunofluorescence staining. The mean ± SD myoblast:fibroblast ratio of the twelve participants was 1.26 ± 1.00 (range: 0.276 to 2.93). We did not detect any differences in muscle stem cell characteristics between cells obtained from females and males (data not shown). We, therefore, combined the data from all muscle cells and correlated the muscle characteristics with individual myoblast:fibroblast ratios. We observed no correlations regarding myoblast:fibroblast ratio and the total number of skeletal muscle stem cells (combined number of myoblasts, fibroblasts and other stem cells) migrating into the artificial wound within all three segments combined at 24 h (R^2^=0.20, F_1,10_=2.56, P=0.141) or 48 h (R^2^=0.02, F_1,10_=0.19, P=0.671) after the scratch assay.

However, there was an inverse correlation between myoblast:fibroblast ratio and migration dynamics for the 12 participants (Figure 3B). Muscle stem cells with a low myoblast:fibroblast ratio demonstrated more cells in the inner segment than to the outer segment compared to muscle stem cells with high myoblast:fibroblast ratio at 24 h (R^2^=0.49, F_1,10_=9.53, P=0.011) and with a non-significant trend at 48 h (R^2^=0.30, F_1,10_=4.33, P=0.064) after the artificial wound healing assay reinforcing the preliminary *in vitro* results from our laboratory

**Figure 3.**
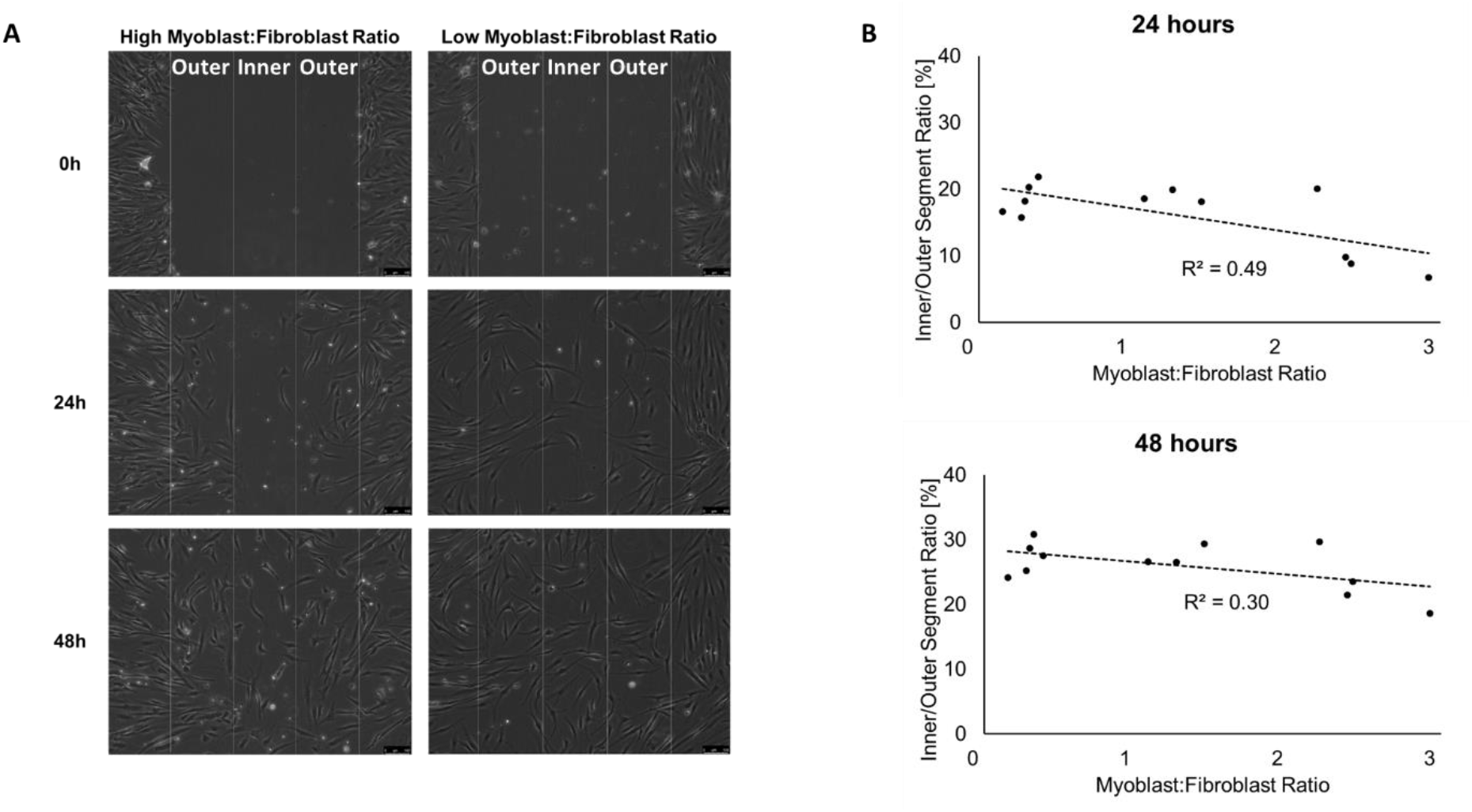
**(A)** Representative images for cell migration of muscle cells with a high myoblast:fibroblast ratio (2.4; left) and with a low percentage of myoblasts (0.3; right) into the artificial wound. The wound area is about 900 μm in width and split into 3 × 300 μm segments (one inner and two outer segments). Magnification is x 10.5, and scale bar is 100 μm. **(B)** Inverse correlations between the myoblast:fibroblast ratio and the migration dynamics of 12 different primary muscle stem cells 24 h (P=0.011) and a trend 48 h (P=0.064) after the artificial wound healing assay.

### Comparison of the Muscle Response between the Repeated Maximal Sprint Protocol and the Muscle Stem Cell Study

In order to determine whether skeletal muscle stem cell ratio (i.e. myoblast:fibroblast ratio) played a role in muscle strength recovery *in vivo*, further studies were performed. Previous investigations have shown that skeletal muscles of different origin, but with similar physiological functions and fibre type composition demonstrate similar transcriptome expression patterns of up to 99% ^26,27^. Further, all limb muscles arise developmentally from the ventrolateral dermomyotome of the segmented paraxial mesoderm ^28^ and these muscles with a similar fibre type composition show low intra-subject ^29,30^ but high inter-subject variability ^30,31^regarding the total amount of stem cells within these muscle fibres. Thus, muscle biopsies were obtained from the vastus lateralis of six participants, who also performed the repeated maximal sprint intervention (they volunteered to provide a muscle biopsy at least three weeks before completing the repeated maximal sprint intervention). As muscle fibre-type composition is similar between the quadriceps and hamstrings ^26^, the stem cell composition of the vastus lateralis was considered representative of both the quadriceps and hamstring muscle groups. The mean ± SD myoblast:fibroblast ratio of the six participants was 1.46 ± 1.06 (range: 0.299 to 2.93). There was an inverse correlation between myoblast:fibroblast ratio and the percentage change in relative hamstring MVC torque measured PRE-to-POST48 *in vivo* (R= −0.945, F_1,4_=33.73, P=0.004; Figure 4A). Thus, participants with a high myoblast:fibroblast ratio showed a delayed hamstring strength recovery 48 h after repeated maximal sprints compared to those with a low myoblast:fibroblast ratio. Further, there was an inverse correlation between myoblast:fibroblast ratio and relative hamstring MVC torque measured POST-to-POST48 (R=-0.943, F_1,4_=17.08, P=0.014; Figure 4B). No correlations were found between the myoblast:fibroblast ratio and changes in quadriceps MVC torque or with any other muscle damage and fatigue biomarker following repeated maximal sprints. These inverse correlations could be confounded by the model assumptions and the relative low participant number. However, post hoc power calculations demonstrate that the bivariate correlation had adequate power (0.97).

**Figure 4.**
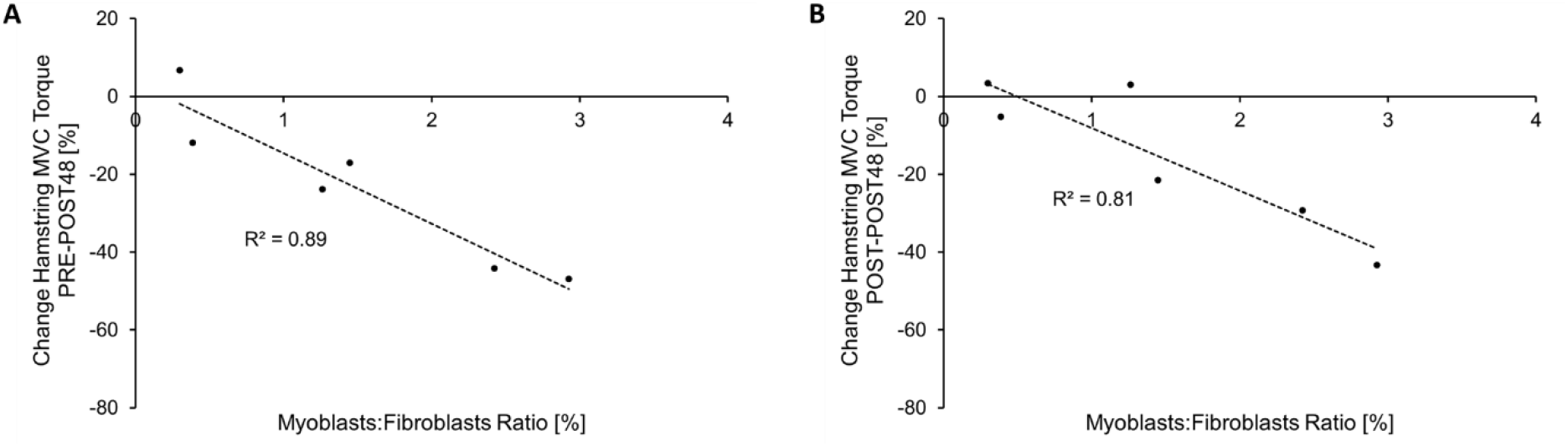
Inverse correlation between the myoblast:fibroblast ratio, assessed in the current in vitro study and the change of hamstring MVC torque measured **(A)** before and 48 h after (P=0.004), and **(B)** measured immediately after and 48 h after (P=0.014) (B) the repeated maximal sprint intervention.

## DISCUSSION

In this study, we have used an interdisciplinary approach to investigate the potential biomechanical, physiological and cellular factors underpinning neuromuscular fatigue following repeated maximal sprints. We have shown that immediate strength loss was associated with reduced hamstring sEMG activity (indicating impaired hamstring motor unit recruitment) and markers of peripheral fatigue, but the magnitude and sustained changes in MVC torque over time (especially in the hamstrings) was largely associated with indicators of peripheral fatigue. Muscle damage biomarkers indicated that the hamstring peripheral fatigue might have been caused predominantly by ultrastructural damage within the muscle tissue. Further, both central and peripheral fatigue caused by repeated maximal sprints appeared to affect the neuromuscular control of running patterns, while a larger BF_LH_ PCSA was related to attenuated hamstring strength loss immediately after the repeated maximal sprint intervention. Finally, our results suggest that a high myoblast:fibroblast ratio leads to a delayed wound closure *in vitro* and to a delayed MVC torque recovery following repeated maximal sprints *in vivo* within the first 48 h, indicating that stem cells of the non-contractile muscle tissue might positively affect the response to muscle damaging exercise.

### Fatigue and Muscle Damage Following Repeated Maximal Sprints

We showed a decreased activity of normalised BF_LH_ sEMG activity immediately after the repeated maximal sprint intervention, but no significant changes in neuromuscular activation using the interpolated twitch technique. The discrepancy between these two methods might be explained by the fact that voluntary activation measured via the interpolated twitch technique investigates all of the hamstring muscles, whilst the normalised EMG analysis was confined solely to the BF_LH_, which is in line with a previous study ^5^. The BF_LH_ might fatigue to a greater degree immediately after repeated maximal sprint related interventions compared to the other hamstring muscles. However, we also provide evidence for peripheral fatigue occurring immediately after the repeated maximal sprints and a delayed recovery at higher frequencies (30-50 Hz) after observing a right shift in the torque-frequency relationship. This may be due to ultrastructural damage predominantly in fast-twitch (which fire at rates from 30-50 Hz) compared to slow-twitch muscle fibres (discharge rates 10-25 Hz) ^32,33^, leading to impaired force generation rather than simply fatiguing the muscle fibres.

Both the quadriceps and hamstring muscle groups showed similar strength loss immediately after the repeated maximal sprints, but the hamstring muscle group showed further strength loss 48 h later compared to the quadriceps. Other studies did not show this additional strength loss for the hamstring muscle group POST48. Differences in the training status of the participants ^7,20^ and in the methodological approaches ^34^ might partly explain the different outcomes. The peak BF_LH_ EMG activity occurs at a more extended knee angle during hamstring isokinetic muscle contraction compared to the peak EMG occurring at a more flexed knee angle for the other hamstring muscles, such as the semitendinosus ^35^. Therefore, we suggest that the BF_LH_ is the key hamstring muscle responsible for decelerating the shaft at the end of the late swing phase. After repeated bouts of high-speed running, the semitendinosus might fatigue prematurely ^36^ and the BF_LH_ would need to substitute the impaired function of the preceding semitendinosus to decelerate the shaft.

During high-speed running, eccentric contractions occur in the hamstrings during the late swing phase (i.e during deceleration of the knee extension) ^37^ and in the quadriceps during the early/mid swing phase (= deceleration of hip extension) ^38^. In general, the likelihood of sustaining a quadriceps strain injury is significantly lower compared to a hamstring strain injury ^2^. If repeated eccentric contractions are one of the main causes for strain injuries, as indicated by several investigations ^39,40^, then the difference in strength loss between the quadriceps and hamstrings observed in our study might be explained by the different levels of eccentric force generated by the muscles. Muscle damage in the quadriceps muscle presumably occurs during the deceleration phase of sprinting and during the backswing phase, when the quadriceps muscle works eccentrically to decelerate the leg with a flexed knee during the early swing phase of high-speed running ^21,38^. Hamstring strain injury, however, occurs with an almost extended leg during the late swing phase ^37^. This extended lever arm might cause higher eccentric force in the hamstring muscles compared to the shorter lever arm with a flexed knee on the quadriceps muscle. Continuously repeated eccentric contractions with the longer lever arm will potentially induce more muscle damage within the hamstrings in total during e.g. a soccer match, compared to the quadriceps muscle group, which might explain the significant different strength loss POST48 repeated sprints in the current study, and potentially the different injury rates between these two muscle groups.

### Kinematic Analysis

Our *in vivo* intervention caused changes in the running kinematics with reduced knee extension in the late swing phase immediately after the repeated maximal sprints. Reduced hamstring muscle strength due to neuromuscular fatigue might trigger a protective mechanism directly after repeated maximal sprints. Afferent signals from the fatigued and damaged hamstrings might activate the Golgi tendon organ ^4^, thus limiting hamstring muscle fibre strain in an attempt to minimise further muscle damage. These kinematic changes were not evident 48 h after the repeated maximal sprints. However, there was a non-significant tendency for prolonged stride duration during running (P=0.08, data not shown) 48 h later and the percentage change of knee extension in the late swing phase of running correlated with changes in hamstring strength both measured from POST to POST48. This indicates that participants with delayed hamstring strength recovery were still not able to fully control running. As hamstring MVC continued to deteriorate 48 h after repeated maximal sprints but quadriceps MVC started to improve, it could be that lower-limb kinematics in the sagittal plane are controlled by the hamstrings more than the quadriceps. That outcome could also have implications for the underlying mechanisms of knee injuries, such as an anterior cruciate ligament injury ^41^, which would need further investigation. Summarised, ultrastructural damage in the hamstring muscles might lead to decelerated movement patterns over time, which could increase the risk for hamstring strain injury during sprinting ^3^.

### The Role of the Extracellular Matrix on the Muscle Response Following Repeated maximal sprinting

Recent investigations have suggested that hamstring maximum eccentric strength and BF_LH_ fascicle length are predictors of hamstring strain injury ^17^. Further, a fatigued muscle is likely to accentuate the risk of muscle strain ^2^. However, we could not find any correlation between BF_LH_ fascicle length and any biomarker of fatigue but BF_LH_ PCSA correlated inversely with hamstring strength loss from PRE to POST. During the late swing phase of sprinting, the hamstring muscles contract eccentrically to decelerate the shaft and to enhance the subsequent concentring shortening contraction for maximal sprinting by using stored elastic energy from the muscle-tendon unit. In comparison to other conventional muscle-damaging interventions ^42^, this dynamic (stretch-shortening) movement might lead to an additional damage of the hamstring muscle connective tissue structure. Therefore, a larger BF_LH_ PCSA might protect against immediate hamstring MVC loss due to a greater ability to transmit the ground reaction forces laterally (from fibre to fibre) ^43^, which might disperse the force more efficiently to the tendon, while the muscle fibres themselves undergo less strain. Further, a greater BF_LH_ PCSA reflects more fibres aligned in parallel, which would be accompanied by more muscle connective tissue of the extracellular matrix, thus potentially protecting the muscle fibres from excessive damage during eccentric contractions.

The stem cells of the extracellular matrix also demonstrated an important role for muscle strength recovery in the subgroup of participants, as there was a strong inverse correlation between myoblast:fibroblast ratio and hamstring MVC torque recovery POST48. Skeletal muscles with a higher availability of fibroblasts around the area of myotrauma might have a better capacity to reorganise the complex extracellular matrix, thus restoring (lateral) force transmission, which results in a faster recovery of muscle strength after muscle damage. This was in line with the myoblast:fibroblast ratio effect on cellular aspects of muscle regeneration and remodelling assessed in primary muscle stem cells *in vitro*. Muscle stem cells with a low myoblast:fibroblast ratio revealed a faster wound closure (i.e. more cells migrated to the inner part of the artificial injury compared to the outer part), in particular 24 h after performing the scratch assay. These results suggest that a larger abundance of fibroblasts has a positive effect at the beginning of muscle repair. We, therefore, assume that repeated maximal sprints with insufficient recovery of previously fatigued and damaged muscles (where the fatigue and damage response is modulated by the muscle size and stem cell composition, respectively) might augment the risk of muscle strains, as appropriate damage to the muscle connective tissue is thought to differentiate between exercise-induced muscle damage and muscle strains^44,45^.

The practical implications of our study are that a 48 h recovery period following repeated maximal sprinting is insufficient, and might increase hamstring strain injury risk. Furthermore, increasing hamstring PCSA via resistance training is likely to reduce peripheral fatigue following repeated maximal sprinting, thereby reducing hamstring strain injury risk.

### Limitations

The current study observed a relationship between human primary muscle cell type (*in vitro*) and physiological biomarkers of skeletal muscle damage/recovery following strenuous exercise. Further research is necessary to confirm these results with a larger sample size regarding the *in vitro* study. However, given the scarcity of data that have examined human muscle stem cell characteristics in association with muscle damage/recovery *in vivo*, and that most of our results were sufficiently powered, we believe that our study represents an important advancement in our understanding of how skeletal muscle recovers following strenuous exercise. There was no relationship between the myoblast:fibroblast ratio and any physiological variables regarding the quadriceps femoris, from which the muscle biopsies were obtained. It has previously been shown that skeletal muscles of different origin, but with similar physiological functions and fibre type composition, demonstrate similar transcriptome expression patterns of up to 99% ^26,27^. For immnunohistochemistry analysis, the ICC (3, k) of 0.83 (95% CIs: 0.59-0.95) indicates a good reliability for the characterisation and the quantification of myoblasts and fibroblasts. Therefore, it is likely, that the correlation between myoblast:fibroblast ratio and the muscle damage-response of the hamstrings but not the quadriceps muscles is explained by more severe ultrastructural damage in the hamstrings than quadriceps. Furthermore, peripheral fatigue can be caused by metabolic perturbations, such as the depletion of intramuscular glycogen ^11^. Therefore, because we did not control diet throughout the study, it is possible that inter-individual differences in baseline muscle glycogen may have influenced the ability to maintain maximal intensity throughout the sprints. However, participants were instructed to eat and drink similar foods two hours before each laboratory visit, and to avoid strenuous exercise for at least 48 h prior to the testing. Further, participants were given sufficient recovery between sprint repetitions and there was a low decrement in sprint performance, indicating that glycogen depletion was probably only a minor factor.

## CONCLUSION

Repeated maximal sprints induce a greater and more prolonged strength loss in the hamstrings compared to the quadriceps muscles. The immediate loss of hamstring function appears to be due to both central (particularly reduced neuromuscular activation of the biceps femoris long head) and peripheral fatigue, while prolonged hamstring strength loss is predominantly linked to peripheral fatigue. Thigh neuromuscular fatigue following repeated maximal sprints alters hip and knee kinematics during running immediately after the repeated maximal sprints, which may lead to an increased hamstring muscle injury risk. Furthermore, biceps femoris long head PCSA was inversely related to hamstring strength loss immediately after repeated maximal sprinting. This suggests a greater PCSA may help transmit more ground reaction force laterally between muscle fibres, e.g. via the extracellular matrix, thus placing less stress during eccentric contractions on the individual muscle fibres and protecting against muscle damage/fatigue. Furthermore, our results suggest that skeletal muscles with an increased number of fibroblasts might have a better capacity to reorganise the complex extracellular matrix, which results in a faster wound closure after substantial muscle damage.

## MATERIALS AND METHODS

### Participants

#### In vivo repeated maximal sprint intervention

Twenty recreationally active and healthy young men (*mean ± SD*; age 20.3 ± 2.87 years; height 1.79 ± 0.05 m; body mass 75.0 ± 7.89 kg) participated in the repeated maximal sprint intervention.

#### In vitro muscle stem cell component

Eight healthy young male (age 21.25 ± 4.27 years; height 1.77 ± 0.05 m; body mass 73.78 ± 5.68 kg) and four healthy young female (age 25.5 ± 1.29 years; height 1.67 ± 0.08 m; body mass 61.40 ± 2.57 kg) participants provided a biopsy of the vastus lateralis muscle for the *in vitro* muscle stem cell component of this study. Six of the eight males also participated in the repeated maximal sprint intervention at least three weeks after providing a muscle biopsy. Only males were recruited for the repeated maximal sprint intervention, as there is some evidence of sex differences in neuromuscular fatigue ^46^. However, both men and women were recruited for the muscle stem cell component due to there being no reported sex differences in stem cell properties and none within our own pilot studies (data not shown). Prior to starting the study, written informed consent was obtained from each participant and pre-biopsy screening was performed by a physician for those participants who volunteered a muscle biopsy. The study was approved by the Research Ethics Committee of Liverpool John Moores University and complied with the Declaration of Helsinki. Volunteers were physically active but were ineligible to participate if they had performed strength training of the lower limbs within 6 months prior to participation in the study, which was determined during pre-participation screening. Further exclusion criteria were: (i) any lower limb injury in the past 12 months; (ii) age under 18 or above 35 years; and (iii) more than three structured exercise sessions per week.

### Experimental Design of the Repeated Maximal Sprint Intervention *in vivo*

Participants were required to visit the temperature-controlled (22-24°C) laboratory on three occasions: (i) familiarisation, (ii) testing day including the PRE and POST assessments; and (iii) POST48 assessments after the repeated maximal sprint intervention. One week prior to the testing day, participants were familiarised with the assessments as well as with repeated maximal sprints (by performing 2-3 submaximal sprints) and BF_LH_ architecture of the hamstring muscle group was assessed via ultrasound. On the test day, participants performed the repeated maximal sprint intervention of 15 × 30 m sprints to induce neuromuscular fatigue/damage in both the quadriceps femoris and hamstring muscle groups. All tests were performed at the same time of the day for each participant. Further, participants were instructed to maintain their normal routine (including eating habits), to refrain from drinking alcohol and to avoid any strenuous exercise 48 h prior to testing and throughout the study, and to refrain from consuming caffeine on testing days. Nothing was consumed throughout the testing sessions except water, which was available ad libitum.

The test battery was always performed in the same order with the right leg of each participant and comprised (i) venous blood sampling [for analysing serum interleukin-6 (IL-6) concentration and creatine kinase (CK) activity]; (ii) hamstrings and quadriceps muscle soreness via visual analogue scale; (iii) isometric maximum voluntary contraction (MVC) torque of the quadriceps, as well as both voluntary and involuntary muscle activation and torque-frequency relationship via electrical stimulation] MVC torque of the hamstring together with normalised BF_LH_ sEMG (see below); and (iv) treadmill running (4.17 m s^−1^) kinematics of the right leg (via an eight-camera motion capture system) PRE and POST48 following the repeated maximal sprint intervention. At POST, kinematic assessments were performed first followed by the aforementioned order of the assessment for practical reasons.

### Maximal Repeated Sprint Protocol

Many athletes in team sports are required to perform repeated maximal sprints, which are characterised by short-duration (<10 s) and relatively longer recovery times (>60 s) between maximal sprint bouts, and have a different physiological demand compared to repeated-sprint exercises with shorter recovery times (<60 s) ^47,48^. Therefore, the repeated maximal sprint intervention consisted of 15 repetitions of 30 m maximal sprints with a deceleration zone of 12 m. The 30 m distance was chosen as the upper average of both the total sprinting distance (346 ± 115 m) of wide-midfielders and the mean recovery time (70.2 ± 25.1 s) between sprint bouts in soccer ^47,49^, which allows the athlete to maintain the performance of the sprint bouts. Similar protocols have been used elsewhere ^6,7,20^. Prior to the repeated maximal sprint intervention, a five-minute warm-up was performed, comprising jogging, dynamic stretching and three self-paced 20 m runs at 60%, 80%, 100% of perceived top speed. During the repeated maximal sprint intervention, the participants were instructed to sprint maximally (verbal encouragement) and to stop within the deceleration zone. Further, they were instructed to move slowly back to the start line and to sit on a chair for the remaining time until the next sprint. The recovery comprised 90 s between repetitions and after every 5^th^ repetition, the participants were allowed to rest for 3 min. Sprinting time during trials was measured and controlled with timing gates (Brower Timing Systems, Draper, UT, USA), which were placed on the start and finish line. Participants started 30 cm before the start line to avoid interfering with timing gates with the arms upon initial acceleration ^21^. Further, heart rate (Polar Oy, Kempele, Finland) and rating of perceived exertion were recorded before and after each repetition. Participants were instructed to wear the same footwear for each testing day. As the fastest sprint time does not necessarily need to occur at the first of the 15 sprint bouts and there was an upsurge in speed of the final sprint, fatigue was assessed with the performance decrement score using the following formula ^19^:

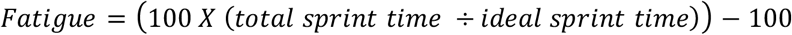

Where total sprint time = sum of time from all 15 sprints; and ideal sprint time = total number of sprints (15) x fastest repetition sprint time. The calculation of this decrement score was also used to quantify changes in heart rate and rating of perceived exertion during the repeated maximal sprint intervention.

### Maximal Voluntary Contraction (MVC)

Three test sessions were conducted with an isokinetic dynamometer (Humac Norm, CSMI Solutions, Massachusetts, USA). As *isokinetic* maximum voluntary contractions (MVC) torque tests are only weak predictors of hamstring strain injury ^50^, we decided to focus on *isometric* MVC quadriceps and hamstring torque at optimal knee strength angles (optimal torque-joint angle relationship, see below) to avoid further fatiguing the participants. The torque signal was interfaced with an acquisition system (AcqKnowledge, Biopac Systems, Santa Barbara, USA) for analogue-to-digital conversion and sampled at a frequency of 2 kHz. The participant was seated in an upright position and securely fastened with inextensible straps at the chest and waist while the arms were held crossed above the chest. The tibiofemoral epicondyle was aligned with the lever arm rotation axis, and the lever arm shin pad was strapped to the leg, 2 cm above the centre of the lateral malleolus. A Velcro strap secured the distal thigh just above the knee. The hip joint angle was set to 85° (180° = supine position) in order to analyse the knee flexor muscle group at a sprint specific angle associated with the late swing phase of sprinting ^51^. Participants were instructed to maximally extend and flex their leg to measure knee range of motion. Quadriceps MVC was measured at 80° knee flexion (0° = full knee extension), as this is the optimal joint angle for peak quadriceps MVC in healthy young men ^52^. Hamstring MVC was measured at 30° knee flexion based on this being the optimal joint angle for peak hamstring MVC during our pilot work. Published studies during the time of data collection used a similar angle of hamstring MVC torque ^53,54^. This was also in line with sprinting kinematics, demonstrating that maximal hamstring muscle lengths during sprinting occur during the late swing phase when the knee is flexed between 30° and 45° ^16^. Prior to isometric MVC assessments, participants underwent a standardised warm up consisting of 10 submaximal isokinetic leg extensions (60°·s^−1^). Participants then performed three isometric knee extension (quadriceps) and flexion (hamstring) at both joint angles (each MVC lasting 2-3 s), with 60 s rest between MVC of a given muscle group. The highest MVC of the three attempts for each muscle group at each angle was used for subsequent analyses. Throughout the tests, participants received verbal encouragement and biofeedback (MVC outputs) were projected onto a screen in front of the participant.

### Hamstring Muscle Voluntary Activation

To measure hamstring muscle voluntary activation capacity via the interpolated twitch technique, stimulation electrodes (12.5 mm × 7.5 mm self-adhesive electrodes (DJO Global, California, USA) were used. The general procedure has been described elsewhere ^5,52,55^. Briefly, the anode was placed proximal to the popliteal fossa, and the cathode was placed beneath the gluteal fold and slightly medial to avoid activation of the vastus lateralis. Protocols were completed with electrical stimulation pads carefully taped down during the sprinting protocol and were additionally marked on the skin with a permanent marker, to ensure a precise relocation for the POST and POST48 tests. Stimulation was delivered by a high-voltage stimulator (DS7AH; Digitimer Ltd., Welwyn Garden City, United Kingdom), and consisted of a doublet using two 240-V rectangular pulses (200 *μ*s pulse width) with an inter-pulse duration of 10 ms (100 Hz stimulation). During each experimental session, relaxed hamstring muscles were stimulated while participants were fixed in the isokinetic dynamometer with the same setting for knee flexion MVC (85° hip angle, 30 ° knee flexion). The amplitude started with 50 mA to familiarise the participants to the stimulation and was gradually increased in 20 mA increments until a plateau in doublet torque was achieved. We decided to use the individual maximal stimulation (100%) intensity despite the fact that other publications used supramaximal stimulation (110-130%) ^5^ as we experienced lower MVC knee flexion torque output beyond 100%. That individual amplitude (162.0 ± 17.4 mA; range: 130–200 mA) was applied during all maximal contractions in the experimental session.

The maximal doublet stimulation was used two minutes later to elicit resting maximal doublet torque in the resting state (control doublet), followed 2.5 s later by a second (superimposed) doublet during an isometric knee flexion MVC. The superimposed doublet torque was always calculated manually from careful selection and inspection of the respective time periods compared to a normal increase in voluntary torque. Voluntary activation was calculated according to the following equation:

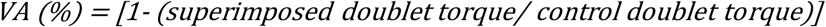

### Surface Electromyography and Antagonist Muscle Co-activation

Surface electromyographic (sEMG) activity was recorded from the vastus lateralis and BF_LH_ to determine the extent of antagonist muscle co-activation during MVCs of the respective muscle group. Previous reports have shown that the vastus lateralis ^56^ and BF_LH_ ^57^ are representative muscles for the quadriceps femoris and hamstring muscle group, respectively. This procedure has been reported in detail elsewhere ^56^. Briefly, once the muscles were identified via palpation, and the skin surface was shaved and cleaned with 70% ethanol, two bipolar Ag-AgCl surface electrodes with an inter-electrode distance of 2 cm (Noraxon duel sEMG electrode, Noraxon, Scottsdale, USA) were placed along the sagittal axis over the muscle belly at 33% of the respective muscle length from the distal end [according to SENIAM guidelines ^58^] and one reference electrode (Ambu Blue, Ambu, Copenhagen, Denmark) was positioned over the medial tibial condyle. The exact location of the electrodes were marked on the participant’s skin with a permanent marker to ensure precise electrode repositioning for the following assessments.

Surface EMG signals were sampled at 2000 Hz (Biopac Systems, Santa Barbara, USA) and then band-pass filtered between 10–500 Hz (AcqKnowledge, Biopac Systems, Santa Barbara, USA). Surface EMG activity of both the agonist and antagonist muscles were analysed by calculating the root mean square of the sEMG signal of a 500-ms epoch around peak MVC. To compare BF_LH_sEMG activity at all three time points, BF_LH_ sEMG of the hamstring MVC at 30° was normalised to the evoked maximum compound muscle action potential (M-wave) of the BF_LH_ (see below). Antagonist muscle co-activation (i.e. quadriceps activation during hamstring MVC at 30° knee flexion, or hamstring activation during quadriceps MVC at 80° knee flexion) was calculated with the following formula (where EMG_max_ is the maximum sEMG of the antagonist muscle when acting as an agonist at the same knee joint angle):

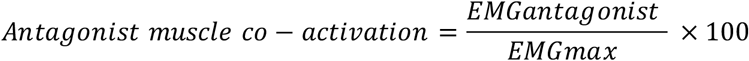

Torque signals, electrical stimuli, and sEMG activity were displayed on a computer screen, interfaced with an acquisition system (AcqKnowledge, Biopac Systems, Santa Barbara, USA) used for analogue-to-digital conversion. Due to technical issues, co-activation data were available for hamstring n = 12; and quadriceps n=10.

### Hamstring Muscle Maximal Compound Muscle Action Potential

The hamstring muscle group was stimulated with single square wave twitch pulses (200 μs duration) using the same electrical stimulator and stimulating electrodes, as described above. While the participant sat resting on the isokinetic dynamometer with the knee angle set at 30° knee flexion, compound muscle action potentials (*M*-waves) were evoked with 10 to 20 mA incremental amplitudes until a maximal *M*-wave (*M*_max_) was achieved. The average amplitude necessary to evoke a maximal *M*-wave was 166.8 ± 19.8 mA; range: 130–210 mA). The maximal *M*-wave was defined as the mean peak-to-peak sEMG response from the three highest observed *M*-waves. Due to inter-individual differences in subcutaneous fat and the (re)location of small sEMG electrodes over a relatively large muscle belly, sEMG amplitude is notoriously variable ^59^. To reduce this inter- and intra-individual variability, we normalised absolute BF_LH_ sEMG to the individual’s BF_LH_ maximal *M*-wave, determined at each testing session ^60^. Due to technical issues, data of BF_LH_ sEMG normalised to maximal *M*-wave was only available for n = 13.

### Torque-frequency Relationship

The torque-frequency relationship was determined by stimulating the hamstring muscle group with single square wave twitch pulses (200 μs duration) at 1, 10, 15, 20, 30, 50 and 100 Hz for 1 s each in a random order and with 15 s rest between each stimulation, using the same electrical stimulator and stimulating electrodes (and location), as described above. The stimulus intensity for 100-Hz stimulation was the amplitude necessary to elicit ~20% knee flexion MVC torque at PRE, and the same amplitude was used for the same test at POST and 48POST. The absolute peak torque at each frequency was normalised to the peak torque at 100 Hz for each time point (PRE, POST and POST48).

### Delayed Onset Muscle Soreness

Using a visual analogue scale that consisted of a 100 mm line (scale 0-10 cm; 0 cm=no soreness; 10 cm= unbearably painful), in conjunction with both a three-repetition bilateral squat (predominantly to determine quadriceps femoris muscle soreness) ^61^ and lunges (predominantly to determine hamstring muscle soreness), participants rated their perceived lower limb muscle soreness along the muscle length immediately after each movement. Muscle soreness was also measured by recording the force required to elicit tenderness at nine fixed sites on the skin over the quadriceps (distal, central and proximal locations of the three superficial quadriceps heads, vastus lateralis, vastus medialis and rectus femoris) and six sites on the hamstrings (distal, central and proximal locations of both BF_LH_ and the medial hamstrings), which were previously marked with a permanent marker to ensure the same measuring position PRE, POST and POST48. At each site, a gradually increasing force was applied by the investigator with an algometer (FPK/FPN Mechanical Algometer, Wagner Instruments, Greenwich, USA) with a maximum of 10 kg/cm^2^. Lying in the prone position with the hip and knee fully extended and muscles relaxed, the participant was asked to indicate when the sensation of pressure changed to discomfort, and the force at that point was recorded.

### Ultrasound

Architectural parameters of the BF_LH_ were assessed using B-mode ultrasound imaging. Participants were in the prone position with the hip and knee fully extended and muscles relaxed. The BF_LH_ was investigated, as this muscle is the most commonly injured hamstring muscle in team sports, particularly during sprinting ^2^. Longitudinal and cross-sectional panoramic ultrasound images of the right BF_LH_ were obtained (Philips EPIQ 7 Ultrasound System, Bothel, USA). The linear transducer (5-18 MHz; aperture 38.9 mm) was carefully placed on the skin with transmission gel and BF_LH_ was scanned (i) longitudinally from its distal (=0% muscle length) to proximal (=100% muscle length) myotendinous junction along a line drawn with a permanent marker to mark the central pathway between the medial and lateral aspects of the muscle (incorporating the intra-muscular aponeurosis (Figure **2**2A); and (ii) cross-sectionally at 20, 40, 60 and 80% along the total muscle length, measured on the skin using a tape measure (Seca, Hamburg, Germany) (Figure 2B).

All images were analysed offline (ImageJ, version 1.51s, National Institutes of Health, Bethesda, USA). Two images for each of the four cross-sectional points were recorded and the image of best quality was used to calculate BF_LH_ muscle volume. The volume of the muscular portion between every two consecutive scans was calculated with the following equation:

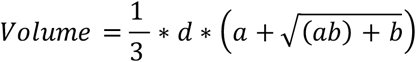

Where *a* and *b* are the anatomical cross-sectional areas of the muscle of two consecutive cross-sectional scans and *d* is the interval distance between the cross-sectional area measurements. The volume of the entire muscle was calculated by summing up all of the inter-scan muscular volumes ^62^. Two full-length sagittal images were then recorded to allow for the measurement of resting BF_LH_ muscle fascicle length and pennation angle, which were both assessed in 3 fascicles at 50% of the total length of BF_LH_. This point was measured offline (ImageJ). A comparison between offline (sagittal ultrasound) and tape measurements of the total BF_LH_ length revealed a very high correlation (R^2^=0.96, P<0.001). Fascicle length was measured by tracing the fascicular path from the upper aponeurosis to the intra-muscular aponeurosis of the BF_LH_. Muscle fascicle pennation angle was determined as the angle between the muscle fascicular paths and their insertion into the intra-muscular aponeurosis. The mean of the three measurements for each parameter were used to determine both fascicle length and pennation angle of the BF_LH_ muscle. PCSA was calculated by dividing BF_LH_ volume by its fascicle length. During the time of data collection, a similar methodological approach was published elsewhere ^63^. One longitudinal image of one participant in the present study was not analysed due to low image quality. Ultrasound scans and image analysis was performed by the same investigator.

### Kinematic and Kinetic Data

Three-dimensional kinematic and kinetic data were synchronously collected at 500 Hz using an eight-camera motion analysis system (Qqus 300+; Qualisys, Gothenburg, Sweden). The data were filtered with a digital dual low-pass Butterworth filter at 20 Hz for motion, as previously described ^64^. Retroreflective markers (12 mm diameter) were placed on anatomical landmarks on the right leg and pelvis, as previously described ^65^. One standing static and two functional motion calibration trials were recorded of the participant PRE, POST and POST48. For the static trial, participants stood with their feet approximately shoulder width apart and knees fully extended. This static trial determined local coordinate systems, the location of joint centres, and the foot, shank, thigh, and pelvis segment lengths of each participant. The functional trials defined functional hip joint centres ^66^ and knee joint axes ^67^. Kinematic data were tracked using Qualisys Track Manager Software (Qualisys). Data processing and analysis were undertaken in Visual3D (C-Motion, Germantown, MD). To examine any changes between the time points, joint angles were normalised relative to the static trial of the accompanying time point for minimising the influence of potential slightly different marker positions between the trials. Lower extremity 3D joint angles and angular velocities were calculated using an X-Y-Z Cardan angle rotation sequence. Investigated variables included peak knee and hip angles, as well as range of motion and time during stance and swing phase of the treadmill run, for all three planes, were calculated as described in previous studies ^64^.

### Motorised Treadmill Run

Participants ran on a motorised treadmill (HP Cosmos Pulsar; Nussdorf, Germany) for 30 s at 4.17 m s^−1^ (0° incline), as high-speed running was of interest. The selected speed was based on pilot testing, which demonstrated that 15 km/h (4.17 m/s) was the fastest speed on a motorized treadmill where the participants still felt comfortable. Motion analysis data were recorded for the last 10 s of the run and data were analysed for at least 6 consecutive strides. Peak knee and hip angle data, for all three planes, were calculated (i) between the initial contact and terminal stance of foot; and (ii) between initial and terminal swing phase. The touchdown of the foot during the treadmill run was determined from the kinematic data as occurring at the local minima and the toe-off during running as the local maxima of the vertical velocity of the head of the fifth metatarsal marker on the foot ^68^.

### Blood Samples

A 10 mL blood sample was drawn from an antecubital vein in the forearm and collected into a serum vacutainer (BD Vacutainer systems, Plymouth, UK). The blood samples were obtained at each time point and left at 22-24°C for 30 min to allow clotting, and then kept on ice when necessary. Serum samples were centrifuged at 1,300 g for 15 min at 4°C. All samples were then aliquoted into 1.5 mL microcentrifuge tubes [Axygen (Corning), New York, USA] and stored at −80°C until subsequent analysis (see below).

### Serum Interleukin-6 (IL-6) Concentration

Serum samples were assayed for IL-6 concentration using commercially available human IL-6 enzyme linked immunosorbent assay kits (Quantikine®, R&D systems, Minneapolis, MN, USA) according to the manufacturer's instructions. The intensity of the colour produced after 20 min was measured with a Thermo Multiskan Spectrum microplate reader (Thermo Fisher Scientific. Waltham, MA. USA) at 450 nm and values were calculated with Excel 365 (Microsoft, v. 365, USA) by generating a four-parameter logistic curve fit. The minimum detectable dose of human IL-6 was 0.70 pg/mL.

### Serum Creatine Kinase Activity

Creatine kinase (CK) activity was assayed using a commercially available CK assay (Catachem Inc., Connecticut, NE, USA), as described in detail elsewhere ^31^. Briefly, 10 μL blood serum were loaded onto a 96-well UV plate. The CK reaction reagent and diluent (Catachem) were prepared as per the manufacturer’s instructions and added to the samples and the change in absorbance monitored continuously over 20 min in a Thermo Multiskan Spectrum plate reader (Thermo Fisher Scientific. Waltham, MA. USA) at a wavelength of 340 nm.

### Capillary Blood Lactate Concentration

Capillary blood samples were taken from the finger-tip via a Safety-Lancet Extra 18G needle (Sarstedt; Nümbrecht, Germany) at rest before and immediately after the repeated maximal sprint intervention. Blood samples were analysed within 60 seconds of collection using a portable blood lactate analyser (Arkray Lactate Pro; Kyoto, Japan).

### Reagents, Chemicals, and Solvents for Muscle Cell Culture *in vitro*

Growth media used for the expansion of human muscle-derived cell populations consisted of Hams F-10 nutrient mix (Lonza, Basel, Switzerland) with added L-glutamine (2.5 mM), 10% heat-inactivated fetal bovine serum (hiFBS; Gibco, Thermo Fisher Scientific, Altincham, UK), 1% penicillin-streptomycin (Life Technologies, Warrington, UK), and 1% L-Glutamine (Gibco). Differentiation media consisted of α-MEM (Lonza), 1% hiFBS, 1% penicillin-streptomycin, and 1% L-glutamine. Phosphate-buffered saline (PBS; Sigma-Aldrich) was used to wash cell monolayers. Desmin polyclonal rabbit anti-human antibody (Cat# ab15200, RRID: AB_301744) was used (1:200) from Abcam (Abcam, Cambridge, UK), and secondary antibody (TRITC polyclonal goat anti-rabbit; Cat# A16101, RRID: AB_2534775) was used (1:200) from Fisher Scientific.

### Muscle Biopsy Procedure

Participants were instructed to avoid any strenuous exercise 48 h prior to the biopsy procedure. Biopsies from the vastus lateralis muscle were obtained under local anaesthesia from each participant, using the Weil-Blakesley conchotome technique as described previously ^69^. The conchotome was inserted through the incision into the muscle belly to obtain the 134 ± 82.7 mg muscle biopsy.

### Extraction of Human Muscle-Derived Cells

The muscle biopsies analysed in this study were isolated and cultured ^31^, as reported previously. Briefly, biopsy samples were transferred with precooled growth media from the muscle biopsy suite to the sterile tissue culture hood (Kojair Biowizard Silverline class II hood; Kojair, Vippula, Finland) within 40 min and muscle biopsy samples were washed three times with ice-cold PBS (0.01 M phosphate buffer, 0.0027 M KCl, and 0.137 M NaCl, pH 7.4, in dH2O). Visible fibrous and fat tissue was removed using sterile scissor and forceps. Samples were cut in small pieces (1 mm^3^) and digested in 5 ml of trypsin-EDTA for 15 min on a magnetic stirring platform at 37°C to dissociate muscle cells. The trypsinisation process was repeated twice. Supernatant derived following each treatment was collected and pooled with hiFBS at a concentration of 10% of the total volume to inhibit further protease activity. Cell supernatant was centrifuged at 450 g for 5 min. Supernatant was discarded and the cell pellet was resuspended in growth media and plated on a T25 cm^2^ culture flask (Corning, Life Sciences, New York, USA) for cell population expansion. Culture flasks were previously coated with a 2 mg/l porcine gelatin solution (90–110 g, Bloom; Sigma-Aldrich, Dorset, UK) to support cell adhesion.

### Expansion of Extracted Cells

The medium was refreshed on the fourth day after the extraction procedure and subsequently every 48 h following two brief washes with PBS. Cells were incubated in a HERAcell 150i CO_2_ Incubator (Thermo Scientific, Cheshire, UK). T25 cm^2^ culture flasks reached 80% confluence after aproximately 10 days and were passaged via trypsinisation. Cells were counted using Trypan Blue exclusion and re-plated on gelatinised T75 cm^2^ culture flasks (Nunc, Roskilde, Denmark). The cells were expanded until passage 3 and then frozen in GM with 10% dimethyl sulfoxide (DMSO) in liquid N2 as a cryopreservant. All experiments were performed on cells between passages 3 and 6 to avoid potential issues of senescence ^25^.

### Characterization of Human Muscle-Derived Cells

The mixed population of human skeletal muscle-derived myoblasts and fibroblasts were characterised by immunofluorescent staining at passage 3 (about 7-10 days after the muscle stem cell isolation procedure before we froze the remaining cells down) for the detection of desmin expressed by myoblasts (desmin positive) and non-myoblasts (desmin negative) to determine the percentage of myoblasts and fibroblasts. Grohmann, et al. ^24^ showed that passaging does not change the percentage of myoblast and fibroblasts and all populations were included for analysis. Previous investigations have determined that the non-myoblast (desmin negative) fraction is highly enriched in fibroblasts, with up to 99% of this fraction being fibroblasts ^70,71^, as our group also observed ^31^. Therefore, non-myoblasts (desmin negative) were referred to here as fibroblasts.

Monolayers were incubated with 25% [vol/vol methanol in Tris-buffered saline (10 mmTris-HCl, pH 7.8, 150 mm NaCl)], 50% and 100% for 5-min to fix the cells and stored at 4°C wet in Tris-buffered saline until further analysis. Fixed monolayers were permeabilised and blocked for 2 h with 5% goat serum and 0.2% Triton X-100 in Tris-buffered saline, prior to staining. Cells were incubated overnight at 4°C with anti-Desmin antibody (1:200). After overnight incubation, the primary antibody was removed, and the cells were washed three times with Tris-buffered saline. Secondary TRITC polyclonal goat anti-rabbit antibody (1:200) was then applied and incubated for 2 h at 4°C. Fluorescent images were captured using live imaging microscopy (Leica DMB 6000; Magnification x 10.5) and analysed via ImageJ cell counter plug-in. A total of four randomly selected areas per well were analysed per individual.

### Wound-Healing Assay, Migration and Differentiation Analysis

For the wound healing assay, 100,000 cells/ml were seeded in gelatinised six-well plates (Nunc, Roskilde, Denmark). Cells were expanded as described above until cell monolayers reached a confluent state, Growth media was removed, monolayers were washed with PBS and cells were damaged by a vertical scrape with a 1-ml pipette tip (width of the wound area, *mean ± S.E.M.*: 896.4 ± 21.24 *μ*m), as previously reported by our group ^31^. PBS was aspirated, damaged cell monolayers were washed twice with PBS to remove cell debris and 2 ml differentiation media was added. Monolayers were imaged with a live imaging microscopy (Leica) for the analysis of cell migration immediately, 24h, and 48h. TIF images were exported from Leica Application Suite and loaded as TIF image stacks in ImageJ with a cell counter plug-in. Cells in the outer and inner segments were then counted (Figure 3A).

Damaged monolayers were imaged at two sites per well in the wound site immediately post-damage (0 h). These image coordinates were tracked and stored to allow subsequent monitoring of the same sites on the wound to reduce this experimental bias. Captured images were exported as TIF image files, and analysed in ImageJ.

### Statistical Analysis

One-way repeated-measures analysis of variances (ANOVA)s were performed to determine whether there was a significant main effect for time (within subject factor) for the following dependent variables: MVC torque, voluntary muscle activation, antagonist muscle co-activation, muscle soreness (for squat lunge via measured via visual analogue scale as well as algometer), rating of perceived exertion, CK activity, IL-6 concentration, and for kinematics data (hip and knee angle parameters). MVC torque data were analysed for interactions and main effects for muscle group and time using two-way mixed design ANOVAs, comparing differences between muscle groups across 3-time points; PRE, POST, and POST48. For within test comparisons, either, independent t-tests, or one-way ANOVAs were used where appropriate. For the torque-frequency relationship, normalised torque at each frequency was analysed using a two-way repeated measures ANOVA, with stimulation frequency (1-100 Hz) and time (PRE, POST and POST48) as the within-groups independent variables. Post-hoc one-way repeated measures ANOVAs were used to determine if the normalised torque at each frequency differed between time points. Bivariate correlations were used to analyse the relation between architectural parameters of the BF_LH_ (volume, fascicle length, fascicle pennation angle and PCSA) and fatigue biomarkers (relative MVC loss normalised to PRE MVC), serum CK activity, serum IL-6 concentration, muscle soreness, knee joint range of motion or changes in range of motion during treadmill running.

Bivariate correlations were used to analyse the relation between myoblast:fibroblast ratio and quadriceps and hamstring MVC, and migration dynamics (total cell migration, cell proportion of inner to outer segment) of the muscle stem cells. Standard guidelines concerning violation of the sphericity assumption to adjust the degree of freedom of the F-test by the Huynh-Felt epsilon if epsilon is greater than 0.75 and to use the more stringent Greenhouse-Geisser adjustment if epsilon is less than 0.75 were followed. Results were expressed as mean ± SD, unless otherwise stated, with statistical significance set at P<0.05. All MVC data were analysed with AcqKnowledge software 4.4 (Biopac-Systems Inc., Goleta, USA) and SPSS 23 Software (IBM Inc., Armonk, NY: IBM Corp) was used for statistical analysis. Occasional missing data are reflected in the reported degrees of freedom.

## GRANTS

This study was supported by a Liverpool John Moores University PhD studentship (P.B.) and the Wellcome Trust Biomedical Vacation Scholarships (R.M.E., M.S.).

## DISCLOSURES

No conflicts of interest, financial or otherwise, are declared by the authors.

## AUTHOR CONTRIBUTION

R.M.E, P.B., B.D., C.E.S., and M.L. conceived and designed the research; P.B., S.T., M.S., M.C., J.A.S., and S.O.S performed the experiments; P.B., S.T. and R.M.E analyzed the data; R.M.E., P.B, M.L. and C.E.S. interpreted the results of the experiments; P.B. prepared the figures; P.B. drafted the manuscript; R.M.E., P.B., S.T., M.S., M.L., B.D., C.E.S., M.C., J.A.S., and S.O.S edited and approved the final version of the manuscript.

## DATA SHARING

The data that support the findings of this study are available from the corresponding author upon reasonable request.

## SUPPLEMENTARY INFORMATION

**Table supplement 4.**
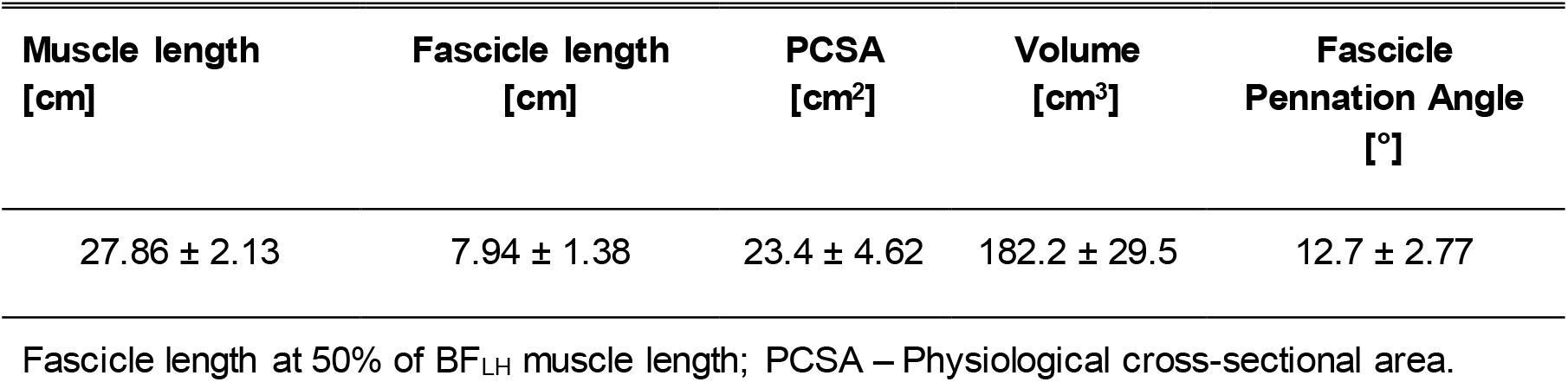
Architectural parameters of biceps femoris long head (mean ± SD).

| Muscle length<br>[cm] | Fascicle length<br>[cm] | PCSA<br>[cm<sup>2</sup>] | Volume<br>[cm<sup>3</sup>] | Fascicle<br>Pennation Angle<br>[°] |
| --- | --- | --- | --- | --- |
| 27.86 ± 2.13 | 7.94 ± 1.38 | 23.4 ± 4.62 | 182.2 ± 29.5 | 12.7 ± 2.77 |

Fascicle length at 50% of BF<sub>LH</sub> muscle length; PCSA – Physiological cross-sectional area.

## Notes

### Competing Interest Statement

The authors have declared no competing interest.

